# Prediction of the Effect of Naturally Occurring Missense Mutations on Cellular *N-Acetyl-Glucosaminidase* Enzymatic Activity

**DOI:** 10.1101/598870

**Authors:** Colby T. Ford, Aneeta Uppal, Conor M. Nodzak, Xinghua Shi

## Abstract

In 2015, the Critical Assessment of Genome Interpretation (CAGI) proposed a challenge to devise a computational method for predicting the phenotypic consequences of genetic variants of a lysosomal hydrolase enzyme known as *α*-N-acetylglucosaminidase (NAGLU). In 2014, the Human Gene Mutation Database released that 153 NAGLU mutations associated with MPS IIIB and 90 of them are missense mutations. The ExAC dataset catalogued 189 missense mutations NAGLU based on exome sequence data from about 60,000 individual and 24 of them are known to be disease associated. Biotechnology company, BioMarin, has quantified the relative functionality of NAGLU for the remaining subset of 165 missense mutations. For this particular challenge, we examined the subset missense mutations within the ExAC dataset and predicted the probability of a given mutation being deleterious and relating this measure to the capacity of enzymatic activity. In doing so, we hoped to learn the degree to which changes in amino acid physicochemical properties are tolerable for NAGLU function.

Amino acid substitution (AAS) prediction methods are mainly based on the sequence and structure information. Simple comparisons between different AAS methods are not only difficult, but also irrational because each method was tested on various datasets and based on varied versions of databases. Currently, several AAS prediction methods have been introduced. PolyPhen-2, an updated version of PolyPhen, is a tool used to predict possible impacts of an amino acid substitution on the structure and function. Users are required to provide protein or SNP identifiers, protein sequences, substitution positions, etc. A score is provided, ranging from 0 to 1, corresponding to the probability of a mutation resulting in no functionality for the enzyme

Once the probability scores were generated, the dataset was then run through multiple machine learning algorithms to generate an applicable model for predicting the enzymatic activity of MPS IIIB-related mutations. This prediction was generated using the PolyPhen-2 probability score and other information about the mutation (amino acid type, location, allele frequency, etc.) as input feature variables. This generated a predicted aggregate score for each mutation, which was then reported back to CAGI. The results of the analysis are significant enough to hold confidence that the scores are decent predictors of enzymatic activity given a mutation in the NAGLU amino acid sequence.

## 1 Introduction

Sanfilippo Syndrome type B disease (Mucopolysaccharidosis IIIB, MPS IIIB) is an autosomal recessive, neurodegenerative disease that primarily afflicts children. Early manifested clinical symptoms includes intellectual disability, behavioral disturbance and death in the second or third decade. MPS IIIB is the inborn error of glycosaminoglycan metabolism and is characterized by the systematic accumulation of substrate heparin sulfate leading to cognitive decline [1]. The lysosomal hydrolase *α*-N-acetylglucosaminidase (NAGLU) performs the function of removing terminal *α*-N-acetyl-glucosaminidase residues from heparin sulfate and lacking NAGLU is one of the four systematic defects that cause MPS IIIB.

The National Institute of Neurological Disorders and Strokes estimates Sanfilippo syndrome type IIIB has a prevalence of 1 in every 200,000 live births [2]. According to the work done by Meikle, et al. in Prevalence of Lysosomal Storage Disorders [3], Sanfilippo syndrome type IIIB generally manifests in young children, carried by an autosomal recessive gene. For those affected, the symptoms are debilitating.

The National MPS Society reports common symptoms include, delayed development and behavioral issues. These symptoms are generally seen in children. As the disease progresses with age, behavioral and cognitive symptoms such as chewing on the hands, difficulties with toilet training, learning disabilities, etc. continue to grow. Other symptoms may include: nausea, digestive problems, and diarrhea as well as the apparent facial formation and distinctive look of MPS-III patients [4].

Currently, there is no known cure for this disease and those diagnosed die at a very early age [4]. An assessment of mutational severity can be time-consuming and costly when one employs traditional wet-lab experiments. In recent years, there has been a drive towards concerted efforts to boost the efficiency of gene mutation studies by implementing computational methods to predict a functional outcome [5]. The effectiveness in characterizing and predicting functional outcomes has proven that the use of computationally-driven predictive models to evaluate the effect of a genetic mutation can save time and money. To increase the use and accuracy of these predictive models, they need to first be utilized and executed in known cases. Continuous research efforts to understand this disease will hopefully lead to a better diagnosis and treatment options. Patients suffering from MPS-III may benefit from a computational framework to predict functional outcomes of particular variants where a more targeted treatment strategy may be developed.

## 2 CAGI

The Critical Assessment of Genome Interpretation (CAGI) provides a unique platform for researchers in bioinformatics to partake in the application of machine learning methods to sharpen the understanding of a life-altering critical disease. The NAGLU use case was part of series IV of the CAGI challenges. The CAGI platform provided the mutation data for MPS-IIIB, which we used in this study.

Results predicting the effects of the NAGLU enzymatic activity were submitted to the Critical Assessment of Genome Interpretation Contest [6]. The submitted prediction is numerically valued in the table 1.

**Table 1:** Prediction scoring table

|  |  |
| --- | --- |
| from 0 | no enzymatic activity |
| to 1 | wild-type level of activity |

**Table 2:** Sample of the CAGI NAGLU dataset for prediction.

| chromosome | variant_position | cDNA_nucleotide_change | AA_substitution |
| --- | --- | --- | --- |
| 17 | 40688337 | 47C>T | A16V |
| 17 | 40688640 | 350T>C | V117A |
| 17 | 40688642 | 352C>T | P118S |
| 17 | 40689424 | 392A>C | Y131S |
| 17 | 40689436 | 404T>G | V135G |
| 17 | 40689442 | 410C>T | T137M |
| 17 | 40689453 | 421T>A | S141T |
| 17 | 40689487 | 455G>A | R152Q |
| 17 | 40689493 | 461T>C | I154T |
| 17 | 40689504 | 472G>T | A158S |
| 17 | 40689538 | 506G>A | S169N |

Included with the above values, amino acid mutation scores and the standard deviation of the predictions were also reported. As shown in Figure 1, these values range from 0.0 to 1.0 and correspond to the interpretations laid out by Table 1. CAGI allowed for a comment to be added in report, including information on whether the mutation was neutral, benign, damaging, deleterious, synonymous or not.

**Figure 1:**
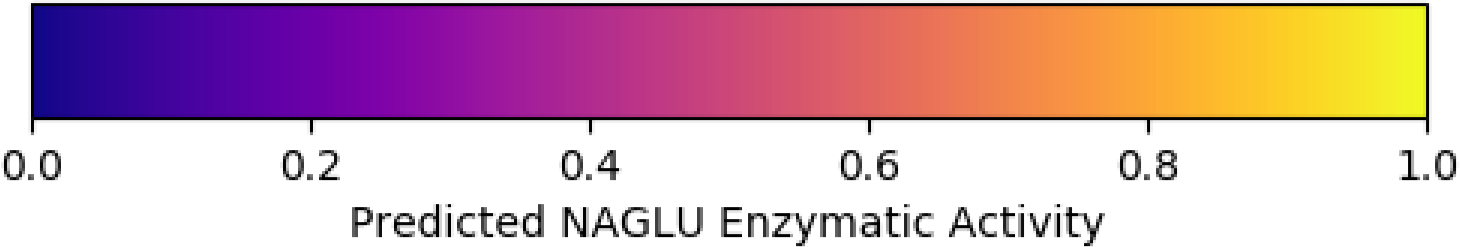
Prediction scores as a function of NAGLU enzymatic activity

**Figure 2:**
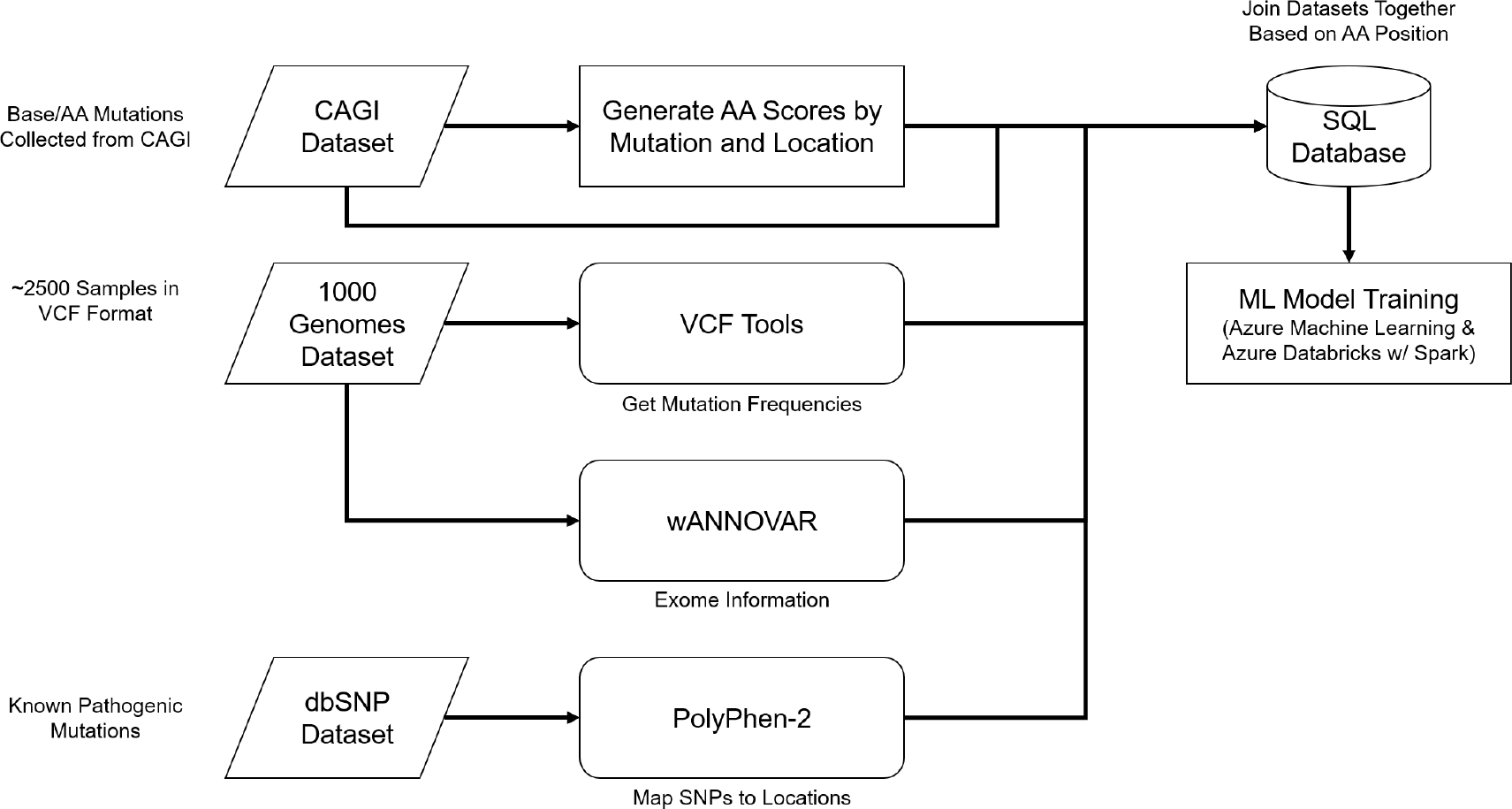
Analysis flow of the data transformations and model generation.

**Figure 3:**
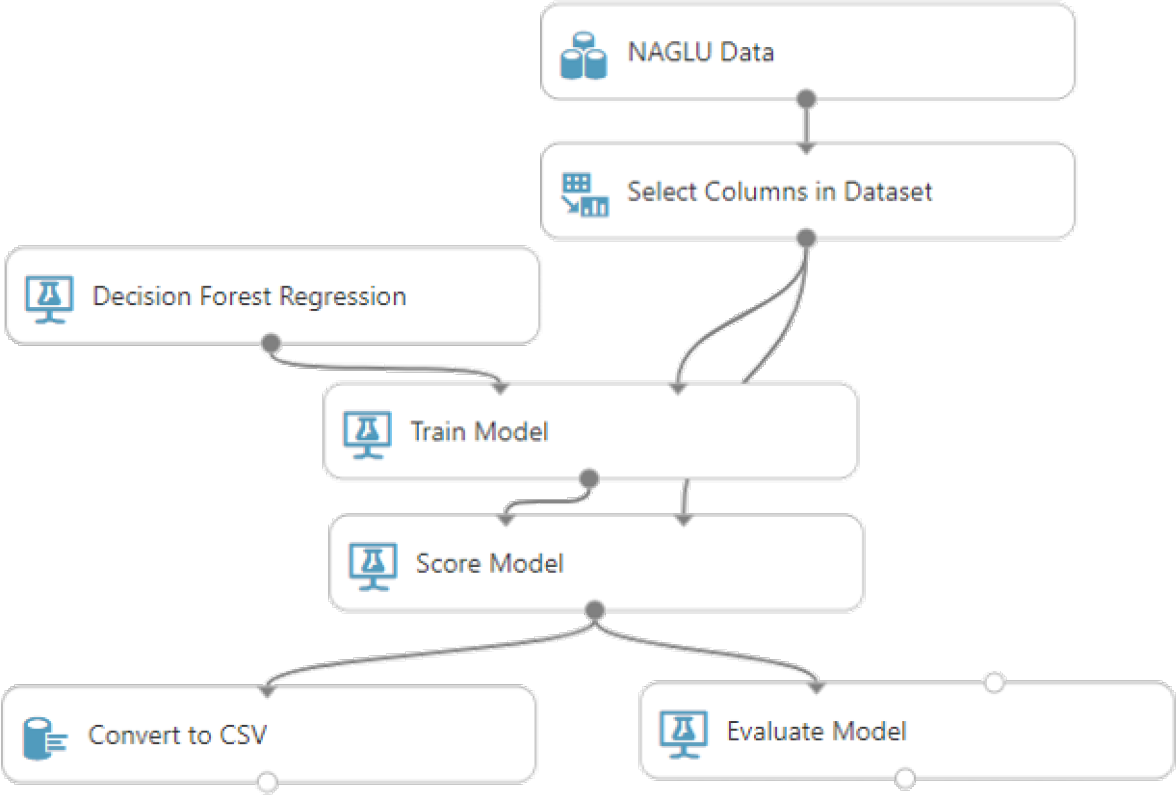
Azure Machine Learning Studio experiment flow.

## 3 Methods

### 3.1 VCFTools

Data from 1000 Genomes is reported in the Variant Call Format (.vcf) [7]. For MPS-IIIB, the interest is only in a particular region on the 17th chromosome. However, only data for the entire 17th chromosome is available for direct use. Once the data was downloaded, the file needed to be trimmed such that we are only working with the region of interest.

Upon downloading the entire chromosome, the file was noticeably large – ~23GB for the .vcf file. At this size, it was far too large to manage simply by using a spreadsheet program. This format was also initially difficult to use in its current tabular layout as it seemed to have pertinent information, but presented in the wrong way for what we want to do with it. *See Table 3*.

**Table 3:**
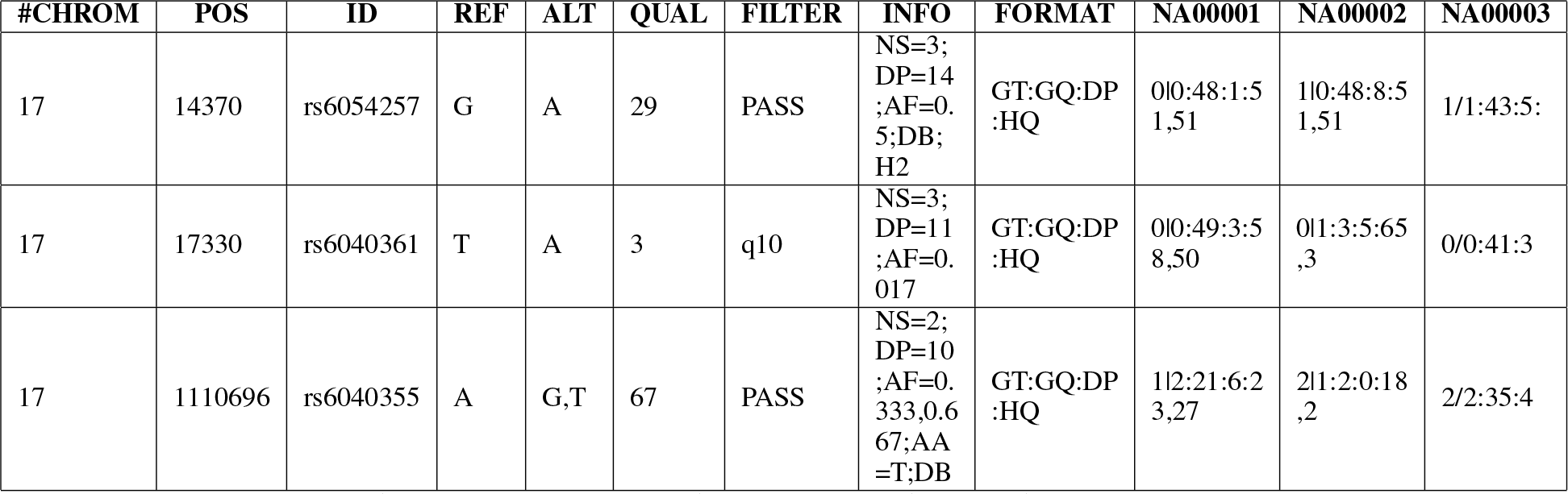
Initial format of the VCF file for human chromosome 17.

Using VCFTools, a Perl add-in for reading and working with .vcf files, we transformed the large input file, shown in Table 3, into a more manageable, tabular format, as shown in Table 4 [8, 9].

**Table 4:**
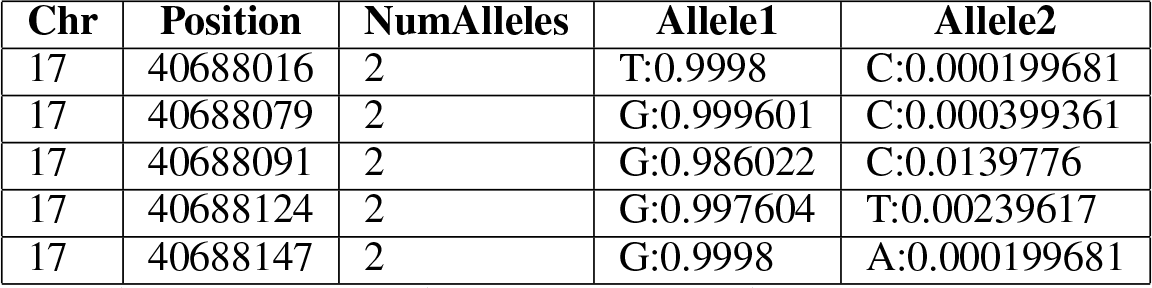
Summarized information after the use of VCFTools

| Chr | Position | NumAlleles | Allele1 | Allele2 |
| --- | --- | --- | --- | --- |
| 17 | 40688016 | 2 | T:0.9998 | C:0.000199681 |
| 17 | 40688079 | 2 | G:0.999601 | C:0.000399361 |
| 17 | 40688091 | 2 | G:0.986022 | C:0.0139776 |
| 17 | 40688124 | 2 | G:0.997604 | T:0.00239617 |
| 17 | 40688147 | 2 | G:0.9998 | A:0.000199681 |

We ran the tool on the entire .vcf file of chromosome 17 and summarized the data to a format that reports the frequency of different alleles at each biallelic site along the chromosome. This information is summarized from the 2,504 unrelated genomes from 26 populations around the globe. By creating a statistical summary and filtering regions of the chromosome not in question, the resulting file size was drastically reduced while providing a relevant indicator of likely pathogenicity based upon incidence.

### 3.2 PolyPhen-2

PolyPhen-2 is a tool that allows for the prediction of functional effects of human nonsynonymous single nucleotide polymorphisms (nsSNPs) [10]. This tool, executed via Perl, takes external references and databases for use in making the functional predictions. We utilized the precomputed *MLC* and *MultiZ* alignments, which are necessary in order to decrease the overall computational time of the program, as well as bundled databases: ncbi-blast-2.2.31+ tools, UniRef100 nonredundant protein sequence databases, and the PDB and DSSP databases.

The tools within PolyPhen-2 run in a sequential order, beginning with the optional first tool, Map SNPs (*mapsnps.pl*). For the CAGI dataset, which provided the reference amino acids and their mutated variant given the position, the PolyPhen-2 input variant format was chosen as the method for running the data through the pipeline. PolyPhen-2 is capable of making predictions based upon two models, *HumDiv* and *HumVar*. For this task, the *HumVar*-model was implemented as the training set used included nsSNPs with minor allele frequencies greater than 1.0%, as well as known disease-associated mutations. Map SNPs is an annotation tool that uses the genomic coordinates for each SNP and produces an output file of those variants which produce missense mutations. This was formatted to be piped into the PolyPhen-2 run (*run_pph.pl*).

As a safety check on the performance of PolyPhen-2, we collected known disease-associated SNPs from the Online Mendelian Inheritance in Man (OMIM) database as a positive control [11]. We also tested the performance on polymorphisms curated by dbSNP as having no associated disease to serve as a negative control set [12]. From this evaluation we felt confident in PolyPhen-2’s ability to reliably calculate the probability of a mutation being deleterious.

For the 1000 Genomes files, the only information present is the genomic position and the variants, making the initial Map SNPs tool necessary in order to extract the nsSNPs. This leads into the next tools for protein variant annotation and probabilistic classification, which includes the initial PolyPhen-2 run (*run_pph.pl*) and Weka prediction (*run_weka.pl*). The PolyPhen-2 output file for SNPs belonging to our CAGI dataset was subsampled and reformatted using Perl in order to highlight columns of interest. A high probability close to 1.0 indicates amino acid change likely to damage the functionality of the protein. PolyPhen-2 generates a prediction output, which we parsed into a tabular format and removed unnecessary attributes, as shown below in Table 5.

**Table 5:** Sample parsed PolyPhen-2 output of CAGI dataset.

| #o_acc | pos | aa1 | aa2 | prediction | pph2_class | pph2_prob | pph2_FPR | pph2_TPR | pph2_FDR |
| --- | --- | --- | --- | --- | --- | --- | --- | --- | --- |
| P54802 | 16 | A | V | unknown | none |  |  |  |  |
| P54802 | 117 | V | A | possibly damaging | deleterious | 0.919 | 0.0594 | 0.811 | 0.0903 |
| P54802 | 118 | P | S | possibly damaging | deleterious | 0.553 | 0.0945 | 0.878 | 0.127 |
| P54802 | 131 | Y | S | probably damaging | deleterious | 1 | 0.00026 | 0.00018 | 0.0109 |
| P54802 | 135 | V | G | benign | neutral | 0.342 | 0.11 | 0.9 | 0.142 |
| P54802 | 137 | T | M | probably damaging | deleterious | 1 | 0.00026 | 0.00018 | 0.0109 |
| P54802 | 141 | S | T | probably damaging | deleterious | 0.974 | 0.0438 | 0.763 | 0.0722 |
| P54802 | 152 | R | Q | benign | neutral | 0.195 | 0.126 | 0.916 | 0.157 |
| P54802 | 154 | I | T | probably damaging | deleterious | 0.993 | 0.0301 | 0.696 | 0.0553 |
| P54802 | 158 | A | S | probably damaging | deleterious | 0.973 | 0.044 | 0.765 | 0.0723 |

The data from the 1000 Genomes is provided as a .vcf file, which provides the genomic coordinates of each of the mutations, but lacks any clear information to ascertain which results are nonsynonymous mutations within the exonic regions for NAGLU. Polyphen-2’s Map SNPs functionality (*mapsnps.pl*) was used to handle this task. However, the output from this tool showed that there were no non-synonymous mutations of the 253 variants called within the region of the NAGLU gene. This result seems unlikely given the large number of variants. A test was done by joining the table of 1000 Genomes SNPs on the table of CAGI nsSNPs, which presented some matches. To determine which variants from the 1000 Genomes data were nsSNPs, and thus variants that could serve as input for PolyPhen-2 and Weka runs, wANNOVAR was used instead. These variants need to be processed through this software in order to make the necessary comparisons across datasets.

### 3.3 wANNOVAR

To accomplish the task of accurately describing the 1000 Genomes dataset, wANNOVAR was employed [13]. wANNOVAR allows for the selection of the hg19 genome build used by 1000 Genomes, and in several different file formats. The input file format is similar to the “genomic position” format used by PolyPhen-2, and it includes the variant genomic position start and end, reference nucleotide, and the variant nucleotide in separate columns. This web-based tool annotates the set of variants, allowing for the extraction of those SNPs designated as being exonic and resultant in nonsynonymous mutations. In addition, the output includes a PolyPhen-2 score derived from the *HumDiv* model.

### 3.4 Database Construction

Given that the machine learning task requires a single, flattened, tabular dataset as input, we curated a database where we could join the various aforementioned features together. These data were housed in a Microsoft SQL Server 2014 server instance.

Importing the datasets was performed using SQL Server Integration Services, an enterprise-grade Extract, Transform, Load (ETL) software. Once the data was successfully written into the database in their respective tables, the tables were joined together by amino acid sequence position. This combines, for example, the .vcf summary with the CAGI data with the PolyPhen probabilities, resulting in a final SQL view that served as the input dataset for our machine learning analysis. *See* Table 7.

**Table 6:** List of database tables used in this analysis.

| Table | Utilized Contents | Source/Generated By |
| --- | --- | --- |
| 1000GenomesSummary | Alleles and their respective frequencies for each position on the chromosome | 1000 Genomes (Summarized by VCFTools) |
| AARef | Reference amino acids by position for NAGLU. Also contains domain information for the sequence. | NCBI |
| CAGIData | Wild Type/Mutant codon base and amino acid changes by position. (Additional information: amino acid types, domain, and an indicator if the amino acid type changed. | CAGI (Additional information added manually) |
| PolyPhen_CAGIPredictions | Predictions for functionality changes in the amino acid mutations from the CAGI dataset input. | PolyPhen2 |
| wANNOVAR1000Genome_summary | Exonic function for each position. (Nonsynonymous mutations) References 1000 Genomes. | wANNOVAR |

**Table 7:** Sample curated data for use in training the machine learning models.

| allele1 | variant_position | AA_substitution | cDNA_nucleotide_change | AA_reference | allele2 | prediction_category | pph2_class | ea_prob | pph2_FPR | pph2_TPR | pph2_FDR |
| --- | --- | --- | --- | --- | --- | --- | --- | --- | --- | --- | --- |
| C | 40688337 | A16V | 47C>T | A | T | unknown | none | null | null | null | null |
| NULL | 40688640 | V117A | 350T>C | V | NULL | possibly damaging | deleterious | 0.081 | 0.0594 | 0.811 | 0.0903 |
| NULL | 40688642 | P118S | 352C>T | P | NULL | possibly damaging | deleterious | 0.447 | 0.0945 | 0.878 | 0.127 |
| NULL | 40689424 | Y131S | 392A>C | Y | NULL | probably damaging | deleterious | 0 | 0.00026 | 0.00018 | 0.0109 |
| NULL | 40689436 | V135G | 404T>G | V | NULL | benign | neutral | 0.658 | 0.11 | 0.9 | 0.142 |
| C | 40689442 | T137M | 410C>T | T | T | probably damaging | deleterious | 0 | 0.00026 | 0.00018 | 0.0109 |
| T | 40689453 | S141T | 421T>A | S | A | probably damaging | deleterious | 0.026 | 0.0438 | 0.763 | 0.0722 |
| NULL | 40689487 | R152Q | 455G>A | R | NULL | benign | neutral | 0.805 | 0.126 | 0.916 | 0.157 |
| NULL | 40689493 | I154T | 461T>C | I | NULL | probably damaging | deleterious | 0.007 | 0.0301 | 0.696 | 0.0553 |
| G | 40689504 | A158S | 472G>T | A | T | probably damaging | deleterious | 0.027 | 0.044 | 0.765 | 0.0723 |
| NULL | 40689538 | S169N | 506G>A | S | NULL | benign | neutral | 0.996 | 0.408 | 0.975 | 0.328 |
| NULL | 40690367 | A181D | 542C>A | A | NULL | possibly damaging | deleterious | 0.327 | 0.085 | 0.862 | 0.118 |
| G | 40690373 | G183A | 548G>C | G | C | probably damaging | deleterious | 0.018 | 0.0393 | 0.748 | 0.0664 |
| NULL | 40690394 | N190S | 569A>G | N | NULL | benign | neutral | 0.912 | 0.15 | 0.932 | 0.179 |
| NULL | 40690401 | F192L | 576C>A | F | NULL | benign | neutral | 0.985 | 0.209 | 0.956 | 0.229 |
| NULL | 40690418 | F198Y | 593T>A | F | NULL | possibly damaging | deleterious | 0.339 | 0.0864 | 0.864 | 0.119 |
| NULL | 40690429 | G202R | 604G>A | G | NULL | possibly damaging | deleterious | 0.122 | 0.0649 | 0.825 | 0.0963 |
| G | 40690437 | M204I | 612G>T | M | T | benign | neutral | 0.969 | 0.181 | 0.947 | 0.205 |
| G | 40690456 | D211N | 631G>A | D | A | benign | neutral | 0.841 | 0.132 | 0.92 | 0.163 |

### 3.5 Machine Learning

Using the conglomerate dataset (shown in Table 7), we utilized multiple machine learning algorithms to generate five different models. Two platforms were used: Microsoft Azure Machine Learning[14] and Apache Spark[15] on Azure Databricks[16]. Both platforms and the various algorithms within all used the same input dataset and the same features. *See Table 8*.

**Table 8:** List of model input variables

| Variable Type | Variable Name |
| --- | --- |
| Label<br>(Dependent Variable) | pph2_prob |
| Categorical Feature<br>(Independent Variable) | allele1, allele2, variant_position,<br>AA_substitution, cDNA_nucleotide_change,<br>AA_reference, prediction_category, pph2_class |
| Numerical Feature<br>(Independent Variable) | pph2_FPR, pph2_TPR, pph2_FDR |

#### 3.5.1 Microsoft Azure Machine Learning

Using the Microsoft Azure Machine Learning Studio, our preliminary model was generated using a Decision Forest algorithm [17, 18]. The input parameters for the Decision Forest algorithm are in *Table 9*.

**Table 9:** Parameters and resulting RMSE and Correlation (compared to actual enzymatic activity) for each machine learning algorithm.

| Algorithm | Platform | Parameters | RMSE | Correlation |
| --- | --- | --- | --- | --- |
| Decision Forest[18] | Azure Machine Learning Studio | Maximum Depth: 128<br>Number of Random Splits per Node: 256<br>Minimum Number of Samples per Node: 1<br>Number of Trees: 10000 | 0.347517 | 0.406035 |
| Logistic Regression[20] | Spark MLlib | Regularization ($\lambda_1$): [0.001, 0.01, 0.1, 0.2, 0.5, 1.0, 2.0]<br>Elastic Net ($\lambda_2$): [0.0, 0.1, 0.25, 0.5, 0.75, 1.0]<br>Maximum Iterations: [1, 5, 10, 20, 50, 100] | 0.396275 | 0.531887 |
| Random Forest[21] | Spark MLlib | Maximum Depth: [2, 5, 10, 20, 30]<br>Maximum Bin: [10, 20, 40, 80, 100]<br>Number of Trees: [5, 20, 50, 100, 500, 10000] | 0.395081 | 0.532796 |
| Gradient Boosted Tree[22] | Spark MLlib | Maximum Depth: [2, 5, 10, 20, 30]<br>Maximum Bins: [10, 20, 40, 80, 100] | 0.395381 | 0.531704 |
| Decision Tree[23] | Spark MLlib | Maximum Depth: [2, 5, 10, 20, 30]<br>Maximum Bins: [10, 20, 40, 80, 100] | 0.395257 | 0.531800 |

#### 3.5.2 Spark MLlib

Revisiting the machine learning approach, we also trained four additional models using the Apache Spark Machine Learning Library (MLlib) [19]. While the Microsoft Azure Machine Learning approach was successful in generating a highly predictive model of NAGLU activity, Spark allows for a much more robust sweep of parameters and cross-validation in a distributed, cluster context.

The four MLlib algorithms used to create additional models were: logistic regression[20], random forest[21], gradient boosted tree[22], and decision tree[23]. Each of the four additional models were trained using a grid search through various parameters as well as a 5-fold cross-validation. *See Table 9*.

An additional change was made in the evaluation of model fitness. As the CAGI competition has since published actual (experimental) enzymatic activity for the mutations in the training data, we used this information in the test dataset to evaluate the models. This was not used in the model training, but to compare the prediction to the actual enzymatic activity values. Then, a model for each algorithm was selected based on minimizing the root mean square error (RMSE) between the prediction and the actual enzymatic activity.

### 3.6 Results

### 3.7 Discussion

Here we show that, by combining heterogeneous information such as reference data (around alleles and amino acids) with MPS-IIIB data (from CAGI) and then process the data using using functional analysis (in PolyPhen-2 with wANNOVAR exon data), we can create a useful training dataset for predicting the enzymatic effects of the mutations related to this disease.

Using various machine learning approaches, we successfully created multiple machine learning models. The winning model, in terms of Pearson correlation is the Spark MLlib-based random forest model. However, the Azure Machine Learning-based decision forest model outperformed based on RMSE.

These models, while not perfect, do have decent predictive power for predicting enzymatic activity for MPS-IIIB.

Predictive models such as these will help researchers understand and pinpoint locations where mutations drastically affect the output and function of NAGLU. By understanding the mutations’ effects, more targeted treatments can be developed to mitigate, reverse, or protect against the functional changes in the mutated enzyme.

Future work should also include further model tuning, complete with the addition of other data. It appears that the correlation between the models’ predictions and the actual tested activity is quite positive, but still lacking. This means that there are likely additional factors that have not been captured by the various datasets used in this study. Thus, finding those other factors and adding them to a model should assist in future model performance, creating a more precise predictions.

